# CAT PETR: A Graphical User Interface for Differential Analysis of Phosphorylation and Expression Data

**DOI:** 10.1101/2023.01.04.522665

**Authors:** Keegan Flanagan, Steven Pelech, Yossef Av-Gay, Khanh Dao Duc

## Abstract

Antibody microarray data provides a powerful and high-throughput tool to monitor global changes in cellular response to perturbation or genetic manipulation. However, while collecting such data has become increasingly accessible, a lack of specific computational tools has made their analysis limited. Here we present CAT PETR, a user friendly web application for the differential analysis of expression and phosphorylation data collected via antibody microarrays. Our application addresses the limitations of other GUI based tools by providing various data input options and visualizations. To illustrate its capabilities on real data, we show that CAT PETR both replicates previous findings, and reveals additional insights, using its advanced visualization and statistical options.

## Introduction

Antibody microarrays use immobilized antibody spots to capture specific protein targets from a given sample; allowing for the binding of many different targets to be quantified [1]. The ever growing number of array validated antibodies have made antibody microarrays an increasingly powerful tool for biomarker and drug target discovery [2], protein modification analysis [3], and protein-protein interaction detection [4]. While the advent of new techniques, such as antibody barcoding [5], and data collection services have made antibody microarray data more accessible, computational tools are still missing to properly process and interpret the resulting datasets. Current tools, such as PAWER [6] and the PMD analysis tool [7], are indeed unable to accommodate data from data collection services or multi-treatment experiments, and offer visualization options that are unsuitable for data exploration and publication.

Here we present CAT PETR (Convenient analysis tool for phosphorylation and expression testing in R), an application that seeks to address these current limitations to create a streamlined, all in one tool for exploratory expression and phosphorylation analysis. To do so, we combine flexible differential analysis options, with interactive and customizable visualizations. To demonstrate the use of CAT PETR, we reanalyze previously published antibody microarray data that looks at how *Plasmodium falciparum* infection modulates signalling protein phosphorylation [8]. We create a series of high quality visualizations that both replicate and compliment the previous studies findings. In addition, by utilizing some of the more advanced differential analysis options in CAT PETR, we are able to narrow down the list of proteins with significantly modulated phosphorylation.

## Implementation

CAT PETR is a web application developed in Shiny (available at https://av-gay-ubc.shinyapps.io/CAT-PETR/), that was designed to work with the results provided by antibody microarray assay services, such as Kinexus Bioinformatics Corp’s KAM 1325 service and Full Moon Biosystems’ antibody array assay service (data provided by other microarray services can similarly be integrated in the future). Both companies’ microarrays have been used to quantify protein expression and covalent modification in a wide range of research, such as in studies of the central nervous system [9], cancer [10], and infectious diseases [11]. The data provided by Kinexus or Full Moon Biosystems can be uploaded directly to CAT PETR, where they undergo several pre-processing steps such as background correction, filtering by technical error, and between-sample normalization. Data provided by other microarray services or collected by the user must undergo the pre-processing steps listed above and be converted to the general user data input format before analysis. This general format consists of tables of gene/protein names and associated expression/phosphorylation metrics, with the option, if appropriate, to specify phosphorylation site information.

Once the data have been uploaded, the user can choose to normalize the data using a logarithmic transformation or variance stabilization normalization [12]. Differential analysis can be completed via traditional pairwise t-tests or Cyber-T t-tests. Cyber-T t-test uses an empirical Bayesian variance estimate to offset the uncertainty in variance caused by low sample sizes and increase power [13]. Cyber-T performs well when compared to other empirical Bayesian methods, and has been widely applied to studies which utilized antibody microarrays, DNA microarrays, and next generation sequencing technologies [14]. Regardless of which method is used, the Resulting P-values can be corrected for multiple testing using either the Benjamini Hochberg or Bonferroni method. At the time of this publication, Empirical Bayesian methods are widely accepted as the most statistically robust option for performing differential expression analysis. As such, we recommend the Cyber-T method for most use cases. As new methods are developed and become standards in the field, we hope to integrate them into CAT PETR as additional statistical options. For more details on the methods and user options, see the user manual, available in supplementary file or directly from the application website.

## Results

We next demonstrate the use of CAT PETR by both replicating and expanding upon the results of a study that studied how infection with *Plasmodium falciparum* modulates signaling protein phosphorylation [8]. The associated antibody microarray data consists of normalized signal intensity data collected from uninfected erythrocytes and erythrocytes infected with *P. falciparum* in the ring, trophozoite, and schizont life stages. Before these data were loaded into CAT PETR, cross reactive antibodies and low signal antibodies were filtered out as recommended [8].

### Volcano Plots and Differential Analysis

2 differential analyses were performed using the uninfected and trophozoite stage infected samples, with the results illustrated via volcano plots. Figure 1A shows the results of the first analysis, which duplicates the statistical methods of the original paper. P-values were calculated using unpaired two sample t-tests without multiple testing correction. The proteins labeled in figure 1A are the same proteins that were identified as significantly modulated within the trophozoite table from figure 3 of Adderley et al. (2020) [8]. Therefore, Figure 1A acts as a fitting compliment to the original figure by showing these noteworthy proteins in the context of every protein sampled. Figure 1B visualizes the results from the second analysis, which uses Cyber-T t-tests and the Benjamini Hochberg adjustment. The Cyber-T t-test introduces more power to the analysis, which partially offsets the more stringent requirements for significance introduced by the multiple testing correction. These stringent requirements allow us to have more confidence in our results, and lets us narrow down the list of significantly modulated proteins to a small group which has been labelled in Figure 1B.

**Figure 1:**
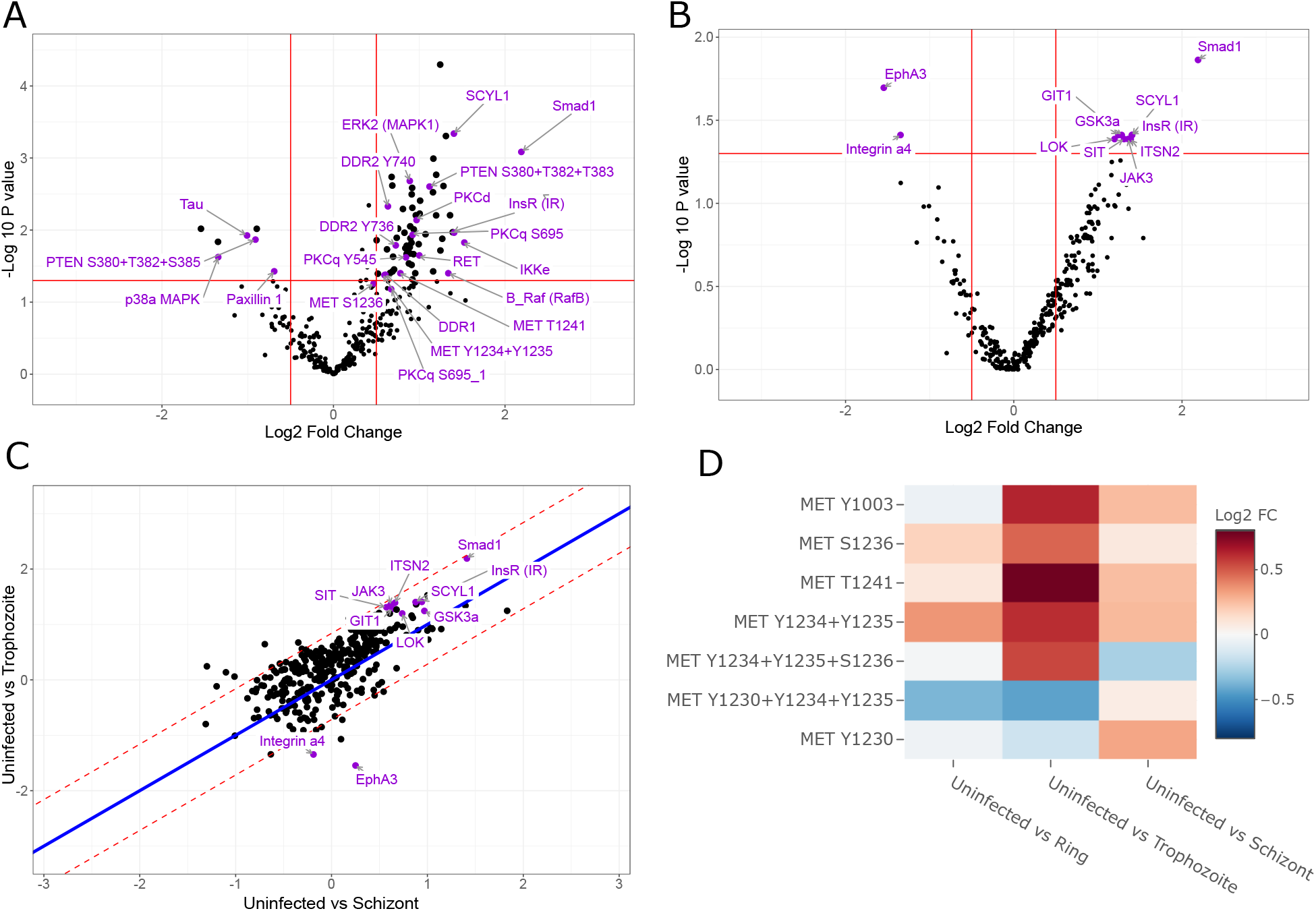
Visualization outputs from CAT PETR (A-B) Volcano plots showing the results of a differential analysis with(B) or without (A) multiple testing correction and Bayesian adjustment. In A, purple labelled proteins are identical to those highlighted in figure 3 of Adderley et al. (2020). In B, labelled proteins are those identified as having significantly modulated phosphorylation. (C) Scatter plot comparing the difference in fold change after infection with trophozoite and schizont stage *P. falciparum*. Purple labelled proteins are the significantly modulated proteins identified in B. (D) Heatmap summarizing the fold changes from all comparisons for MET phosphorylation sites.

### Scatter Plots, Heatmaps, and Treatment Comparisons

CAT PETR provides options for comparing fold changes across multiple control-treatment comparisons using scatter plots and heatmaps. Figure 1C compares fold changes after infection with trophozoite and schizont stage pathogens. We can observe that the 2 most downregulated proteins in the trophozoite stage, identified as significant in figure 1B, return to nearly uninfected levels during the schizont stage. This observation exemplifies the additional insights that can be gained from these scatter plots. Heatmaps provide another opportunity to observe differences in fold change, but for every comparison at once. Figure 1D replicates the results of figure 5A from Adderley et al. (2020) [8], but using a cleaner and more standard format. Proteins/genes can be easily added or to these heatmaps using the app’s searchbar, or by clicking on the points in the other plot types. In addition, these heatmaps have customizable colour palettes, colour scales, and tile sizes to fit into any publication.

## Discussion

Two of the most recent and most extensive antibody microarray data analysis tools include PAWER [6], an R Shiny application, and the PMD analysis tool [7]. Both of these tools can perform differential analysis and provide visualizations, but they also both suffer from several limitations. For example, both apps only accept gpr files as inputs. While gpr files are very commonly used, accepting only one specific file type makes these tools incompatible with many alternative data formats, such as those provided by data collection services. Another limitation is that these apps can only compare 2 treatment groups at a time, making it awkward and time consuming to analyze multi-treatment experiments. Finally, the visualizations provided by these tools are in the form of static PNG images. The lack of interactivity makes these visualizations sub-optimal for data exploration, and the lack of customizability makes them difficult to integrate into publication ready figures. CAT PETR specifically attempts to address the shortcomings of other applications. It provides input options for data from multiple microarray services while also providing input options for generic tables of gene expression data, which allows for CAT PETR to potentially accept data from any file format after appropriate pre-processing. It provides visualization options for working with individual or multiple treatment groups and all visualizations are interactive, customizable, and available to download as editable SVG files.

In conclusion, we have created an easily accessible tool for antibody microarray analysis that addresses the current limitations of similar apps. It includes numerous options for data input types and statistical analysis techniques and has demonstrated effectiveness when applied to real data. It is made available as a web application or can be downloaded onto one’s personal computer via Github. A user manual including a tutorial and detailed instructions for different use cases and example input and output file formats is available as supplementary file, or as part of the application.

## Acknowledgments

We would like to thank J. Adderley and C. Doerig for their assistance in replicating their original statistical methods.

Funding for this work was provided by CIHR PJT-148646 and PJT-152931.

Steven Pelech and his family are the majority shareholders of Kinexus Bioinformatics Corporation.

