## Supplemental file for "CAT PETR: A Graphical User Interface for Differential Analysis of Phosphorylation and Expression Data"

### CAT PETR

December 2022

### 1 Introduction

#### 1.1 What is *CAT PETR*?

*CAT PETR* (**C**onvenient **A**nalysis **T**ool for **P**hosphorylation and **E**xpression **T**esting in **R**) is an R shiny application that provides a user friendly interface for the statistical analysis and visualization of phosphorylation and/or expression data. *CAT PETR* was originally designed for use with the raw text files provided by Kinexus' KAM 1325 antibody microarray service. The latest version now uses a general input format that allows for expression and phosphorylation data collected using any method to be analyzed. differential analysis in *CAT PETR* is performed via either traditional t-tests or Cyber-T t-tests [1] which use an empirical Bayesian variance estimator to adjust for low sample sizes. In addition, *CAT PETR* allows users to explore their results via interactive volcano plots, scatterplots, and heatmaps.

#### 1.2 Development and Requirements

*CAT PETR* has been developed using R 4.2.1 and Shiny 1.7.2. It uses many different R packages such as Tidyverse, purrr, plotly and shinyWidgets. In addition, *CAT PETR* uses the source code provided by the makers of the Cyber-T web-server [1] in order to perform t-tests with empirical Bayesian variance estimation. For ease of use, *CAT PETR* is hosted as a web application via https://av-gay-ubc.shinyapps.io/CAT-PETR/.

#### 1.3 Running *CAT PETR* on the web

*CAT PETR* is hosted on https://av-gay-ubc.shinyapps.io/CAT-PETR/. The web version of CAT PETR works on all operating systems and browsers.

#### 1.4 Running *CAT PETR* locally

In some instances, such as a lack of consistent internet, it may be preferable to install *CAT PETR* directly onto a personal computer rather than using the web server version. To install *CAT PETR* locally, first ensure that R and Rstudio are installed (Installation instructions can be found here https://www.r-project.org/ and here https://www.rstudio.com/products/rstudio/download/ respectively). Then, download or clone the *CAT PETR* GitHub repository (https://github.com/Raif-fl/CAT-PETR). Now, open and run the 'Install_Dependencies.R' file in Rstudio. After the dependencies are done installing, open up the app.R file and run *CAT PETR* by hitting the run app button in Rstudio or by running the shiny::runApp() command.

### 2 General Features

*CAT PETR* has two main tabs: a data upload tab which allows the user to upload their own sample data and compare samples via two sample t-tests, and the visualization section which allows the results of the comparisons to be visually analyzed.

#### 2.1 Data Upload

Any analysis performed with *CAT PETR* will begin on the data upload tab. This is the tab where the user uploads the data from each of their samples and defines which samples will be statistically compared to each other. An example of this tab is shown in Figure 1 of the tutorial.

The data upload tab can be split into two sections, a left sidebar and a main panel. The sidebar on the data upload tab has 3 basic inputs:

1. Radiobuttons that allow the user to specify if they are working with general user defined data, data that has already been previously analyzed by *CAT PETR*, data provided by Kinexus Bioinformatics' KAM-1325 microarray service, or the data provided by Full Moon Biosystems' antibody array services.

2. A browse button which allows the user to search their computer for their data files.

3. A download processed data button which allows the user to download the results from each statistical comparison performed.

The main panel contains a console log. This log will display the progress of your statistical analysis along with any warning or error messages that may occur. Below the console log are two buttons. One of them clears the contents of the log while the other cancels whatever analysis is currently running.

##### 2.1.1 Input Type

*CAT PETR* Excepts 4 different kinds of inputs: general user defined data within a set of csv files, Pre-analyzed *CAT PETR* data within a set of csv files, the raw data txt files provided by Kinexus Bioinformatics' KAM-1325 microarray service, and the Microsoft Excel files provided by Full Moon Biosystems' antibody array services.

##### 2.1.2 User Data

- **Data Format:** User data should be in the form of a series .csv files which contain tables of phosphorylation and expression data. It is expected that this data has already gone through pre-processing steps such as background correction, technical replicate averaging, and between sample normalization. The *CAT PETR* input format (shown below) includes 4 different types of columns. A Name column which contains the gene/protein name, a P_site column that contains the name of the targeted phosphorylation site, an identifier column which contains the gene/protein ID, and a Rep column which contains the data from one biological replicate. Of these columns, the only ones that are required are the Name column and at least 1 Rep column.

  Additional Rep columns can be added in the form Rep_N (Rep_1, Rep_2, Rep_3...). The P_site column can be omitted if the data does not relate to phosphorylation. The identifier column can also be omitted if the Name column already acts as a unique identifier.

  Each csv file uploaded to *CAT PETR* should represent all of the data from one treatment group (e.g. control, treated, infected, mutant). The names of the data files will be used for some of the graph titles and labels, so short filenames are recommended.

| Name | P_site | Identifier | Rep_1 |
|---|---|---|---|
| 4E-BP1 | Pan-specific | NN166-2 | 270 |
| 4E-BP1 | T37+T46 | PN550 | 7961 |
| A6 | Y309 | PK501 | 7460 |
| A6r | Y309 | PK502 | 6465 |
| AAK1 | S637 | PK503 | 9641 |

- **Phosphorylation Site Options:** If *CAT PETR* detects that a phosphorylation site (P_site) column is present in the uploaded files, then 2 additional input options will appear as shown in Figure 2 of the tutorial. These inputs include a text input which allows the user to specify the name used for pan-specific phosphorylation data entries, and some radiobuttons that allow the user to specify if they would like to analyze only the pan-specific phosphorylation data or only the phospho-site specific phosphorylation data.

- **Comparison and Analysis Options:** After the data has been uploaded, the choose comparisons button is used to define which of the data files will be compared to each other. This selection is done using a bucket list that allows the user to choose which of the files uploaded will be treated as control and treatment samples. These samples are compared on a one to one basis as shown in the Figure 3 of the tutorial.

  After the comparisons have been chosen, the user is given the option to select a method of variance stabilization normalization to apply to their data. The first option is a natural logarithmic transformation. The second is a variance stabilization normalization (VSN). VSN is an arsinh based transformation that is often preferable over a logarithmic transformation because it can handle values at or below zero. For more information on the Cyber-T method, please see the appendix in section 4 and the VSN reference [2]. Users can choose whether they want to run a traditional t-test or a Cyber-T t-test. The Cyber-T t-test is highly reccomended for experiements with few biological replicates ($< 5$). For more information on the Cyber-T method, please see the appendix in section 4 and the Cyber-T reference [1].

###### 2.1.3 *CAT PETR* data

- **Data Format:** A pre-analyzed set of .csv data files that were downloaded from *CAT PETR* using the download button on the data upload page. As soon as data files of this type are uploaded, the user can move on to the visualization tab as no additional analysis is needed. It is possible to upload .csv files that were not outputted by CAT PETR as pre-processed data. However, this is not recommended as one would need to alter the column names and values from their data until it has a format identical to what is outputted by *CAT PETR*.

###### 2.1.4 Kinexus KAM-1325 data

- **Data Format:** The raw txt files provided by Kinexus can be loaded directly into *CAT PETR* without alteration.

- **Additional Processing:** The steps to Process the KAM-1325 files are nearly identical to those used for the un-processed csv files with the inclusion of one additional step. Immediately after uploading the raw Kinexus files, one additional input appears. This is a slider which allows the user to select the cutoff for the percentage error between technical replicates where the percentage error is calculated as $\frac{\text{replicate intensity} - \text{mean(replicate intensities)}}{\text{mean(replicate intensities)} \times 100}$. This value should be adjusted based on what the user considers an acceptable amount of error between technical replicates. Apart from this extra input, one can proceed with the same steps used for any other user uploaded csv files.

###### 2.1.5 Full Moon Biosystems data

- **Data Format:** The Microsoft Excel files provided by Full Moon biosystems can be loaded directly into *CAT PETR* without alteration.
- **Additional Processing:** As with the files provided by Kinexus, the steps to Process the Full Moon Biosystems files are very similar to those of any data type with the addition of a couple of extra inputs. Immediately after uploading the Full Moon Biosystems' Excel files, 2 additional radio button inputs appear. The first allows the user to choose which type of antibody array service they are using, a phosphorylation array (e.g. Phospho Explorer Array) or an expression array (e.g. Cell Cycle Antibody Array). The second additional input is only visible when working with phosphorylation arrays and allows the user to choose which kind of phosphorylation site specific data to analyze. Full Moon Biosystems' phosphorylation arrays generally use 2 different types of antibody for each phosphorylation site they are testing. The first kind binds to the phosphorylation site regardless of whether or not it is phosphorylated, and is therefore phosphorylation independent. The second kind only binds to the site if it is phosphorylated, and is therefore phosphorylation dependent. After these extra inputs are specified, the user can proceed with the same steps used for the other analysis options.

##### 2.2 Visualization

The visualization tab contains three separate sub-tabs; one for each of the plots created by *CAT PETR*. While each tab does contain unique inputs and visualizations, they all have similar structures. The sidebar panel for each tab can be roughly split into three sections:

1. A set of check boxes that allow the user to specify if the plot labels will include the gene/protein name, phosphorylation site, and identifier.
2. A set of plot specific inputs.
3. Buttons that allows the user to download the data used to create the plot they are currently viewing. For the volcano plot and scatter plot, the user can choose to either download the data for every point on the graph or just the points selected by the gene/protein search bar.

The main panel for each tab can also be split into three rough sections:

1. At the very top of the main panel are the appearance option and plot download buttons which are represented by cog and camera icons respectively. The appearance option

button opens a pop-up menu that provides a variety of inputs which allow the user to alter the sizing, spacing, and color of different plot elements. The download button downloads the currently visible graph as a high quality .svg file.

2. The plot itself.

3. A gene/protein searchbar that allows the user to pick specific genes/proteins to label/view on the plots. Typing in the name or identifier for a gene/protein will bring up a list of similar names and identifiers. There are additional methods for adding genes/proteins to the searchbar which are dependent on the tab you are using, such as clicking on data points or hitting the 'add labelled entries to searchbar' button.

###### 2.2.1 Volcano Plot

The volcano plot tab allows users to easily visualize which genes/proteins were most effected by a treatment. The volcano plot itself allows the user to add additional points to the gene/protein searchbar by clicking on the data points. The slider below the volcano plot allows the user to quickly switch between plots from different control-treatment comparisons. The volcano plot specific input options on the sidebar include:

- A numeric input that allows the user to specify how many of the top effected genes/proteins to label. These top genes/proteins are defined as the ones furthest from the origin point of the graph (0,0) as defined by the euclidean distance. A small switch underneath this input allows for all of the points outside of the log fold change and P-value cutoffs to be labeled.

- A button which adds all of the labeled genes/ proteins to the searchbar. This is useful when one wants to observe how the top effected genes/proteins in one comparison are effected in other comparisons.

- 2 slider bars which control the cutoffs used for the log fold change and log P-values. These will adjust the red dotted lines seen in the volcano plot.

###### 2.2.2 Scatter Plot

The scatter plot tab allows users to easily compare the log-fold changes from separate control-treatment comparisons. This allows for the easy identification of genes/proteins that were effected differently by different treatments. The scatter plot itself has the log fold change from one control-treatment comparison on the X-axis and the log fold change from a different comparison on the Y-axis. The line drawn through the center of the plot represents a one-to-one ratio between the two comparisons while the dotted lines show the quantiles of the data. Clicking on one of the plot points will add that point to the gene/protein searchbar, similarly to the volcano plot. The slider below the scatter plot allows the user to quickly change between different scatter plots. The scatter plot specific input options on the sidebar include:

- A define axes button that allows the user to specify which of the control-treatment log fold changes will be on the X and Y-axes of the scatter plot. Hitting this button will bring up a popup window with bucket lists similar to the ones used to define the control-treatment comparisons on the data upload tab. Dragging and dropping comparisons from the left-most bucket allows the user to select which comparison will be displayed on each axes.

- A numeric input that allows the user to specify how many of the top affected genes/proteins to label. These top genes/proteins are defined as the points with the greatest distance from the one-to-one ratio line (the blue line) shown on the plot. A small switch underneath this input allows for all of the points outside of the quantiles to be labeled.
- A button which adds all of the labeled genes/ proteins to the searchbar.
- A slider which allows the user to specify the width used for the quantiles.

###### 2.2.3 Heatmap

The heatmap tab allows users to view the log fold changes across all of the control-treatment comparisons simultaneously for a select group of proteins. Unlike the other plots, the heatmap only shows data for proteins that have been selected using the gene/protein searchbar. As such, the heatmap will remain blank if the gene/protein searchbar is empty. The heatmap specific sidebar inputs include:

- A series of checkboxes that allow the user to decide which of the control/treatment comparisons will appear as columns on the graph.
- Drop down options that allow the user to specify the type of color bar they are using (diverging or sequential) and the color palette used for the color bar.
- A slider which allows the users to specify the range of the color bar. There are two options for this range. Auto limits automatically sets the maximum and minimum range values to just above and just below the maximum and minimum fold change values of the genes/proteins shown on the heatmap. Manual limits opens up some numeric inputs that allow the user to specify the maximum and minimum range values for themselves.
- A numeric input that allows the user to choose which of the heatmap columns to sort by. The rows will be sorted from largest to smallest fold change based on the chosen column.

### 3 Tutorial

This is a simple tutorial that covers the data upload and visualization sections of CAT PETR when working with protein phosphorylation data. The example data used is antibody microarray data collected from biopsied pancreatic cells from patients who underwent 5 different drug treatments: No treatment (C), treatment with B-IT (T_BIT), treatment with Sivelestat (T_SIV), treatment with Tebipenum (T_TEB), and treatment with Telaprevir (T_TEL).

#### 3.1 Data Upload Tab

1. The example data for this tutorial can be found on the CAT PETR OSF page https://osf.io/qmd8e/. Please save the data from the example user data folder onto your computer in its own folder.

2. Go to the 'Data Upload' tab. The appearance of this tab is shown below.

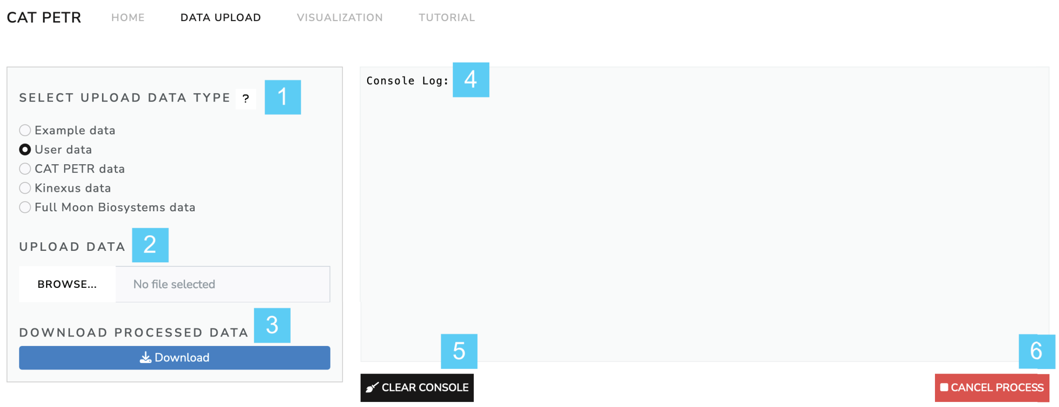

Figure 1: (1) Selection boxes specifying if uploading un-analyzed user data, pre-analyzed CAT PETR data, the raw data files from a Kinexus KAM-1325 microarray , or the Microsoft Excel files provided by Full Moon Biosystems' antibody array services. Question mark brings up example input formats. (2) Data upload section. (3) Button which downloads processed data as a .zip file. (4) Console log which displays the analysis progress along with warnings, errors, and messages. (5) Button which clears the console log. (6) Button which cancels any active analysis processes.

3. Choose the 'User Data' option for the data upload type. Then, click on the browse button and navigate to wherever you have saved the example data. Note that each data file will represent a single sample/treatment group. Use the shift key to select all of the csv files and upload.

4. You will notice that three additional input options appear.

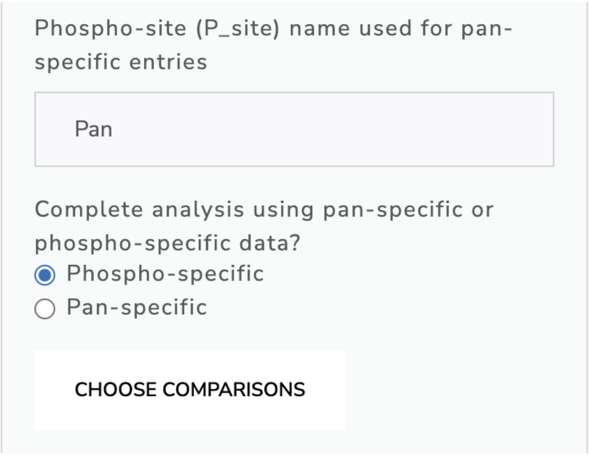

Figure 2: Additional phosphorylation options

The first allows the user to enter what name was used in the Phosphorylation site (P_site) column to specify that the row contains pan-specific phosphorylation data. The second allows the user to clarify if they are interested in analyzing pan-specific data or phospho-site specific data. These two input options will only appear if an optional P_site column is present in the data. For now, leave both inputs as the defaults.

The last new input is the 'choose comparisons' button which will be used to decide which of our samples we want to statistically compare to each other. Click this button and move on to the next step.

5. Clicking the 'choose comparisons' button brings up a modal which contains 3 bucket lists. These lists allow the user to drag and drop their samples into the control or treatment categories. Samples in the control category are compared to samples in the treatment category that are directly adjacent to them as demonstrated in the image below. Please choose controls and treatments to match this image.

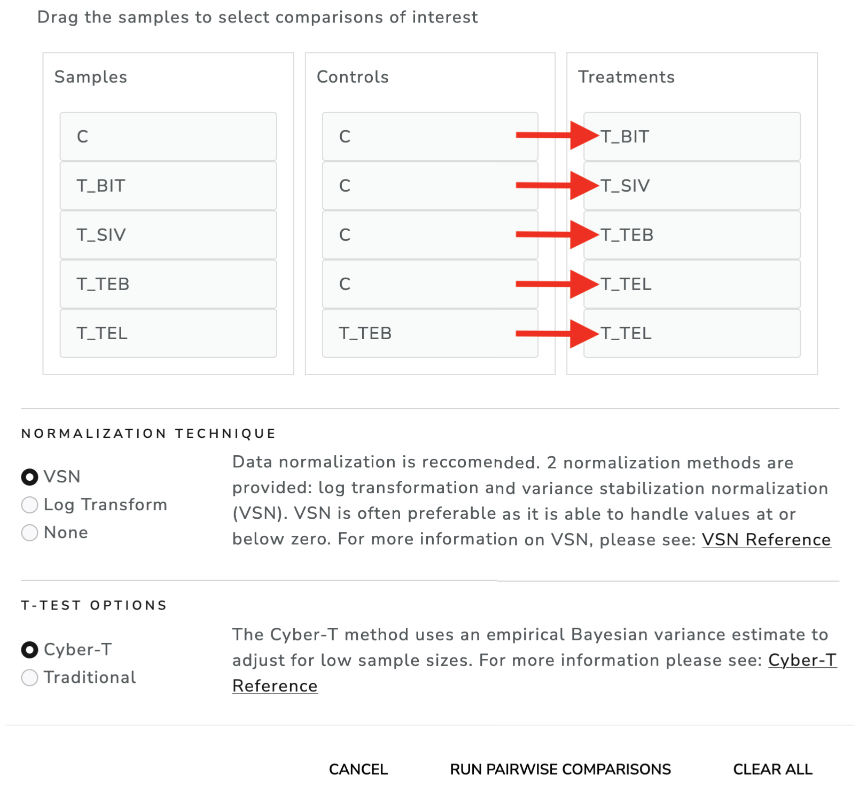

Figure 3: Comparison selection bucket list

6. Once the controls and treatments are selected, the user must the choose normalization and t-test methods for the differential analysis. The normalization techniques offered include Log Transformation and VSN. Please select VSN and Cyber-T and then hit the 'run pairwise comparisons' button. The comparison should be completed in under 30 seconds. Once completed, we can move on to the next section of the tutorial. For more information on VSN and Cyber-T, please see the VSN Reference and the Cyber-T Reference respectively.

#### 3.2 Visualization Tab

The visualization tab contains three sub-tabs: the Volcano Plot tab, the Scatter Plot tab, and the Heatmap tab. We are going to start with the Volcano Plot tab.

##### 3.2.1 Volcano Plot

7. Switch over to the the visualizations tab using the tab-bar at the top of CAT-PETR. The volcano plot will load automatically after you switch tabs and should have the appearance shown below:

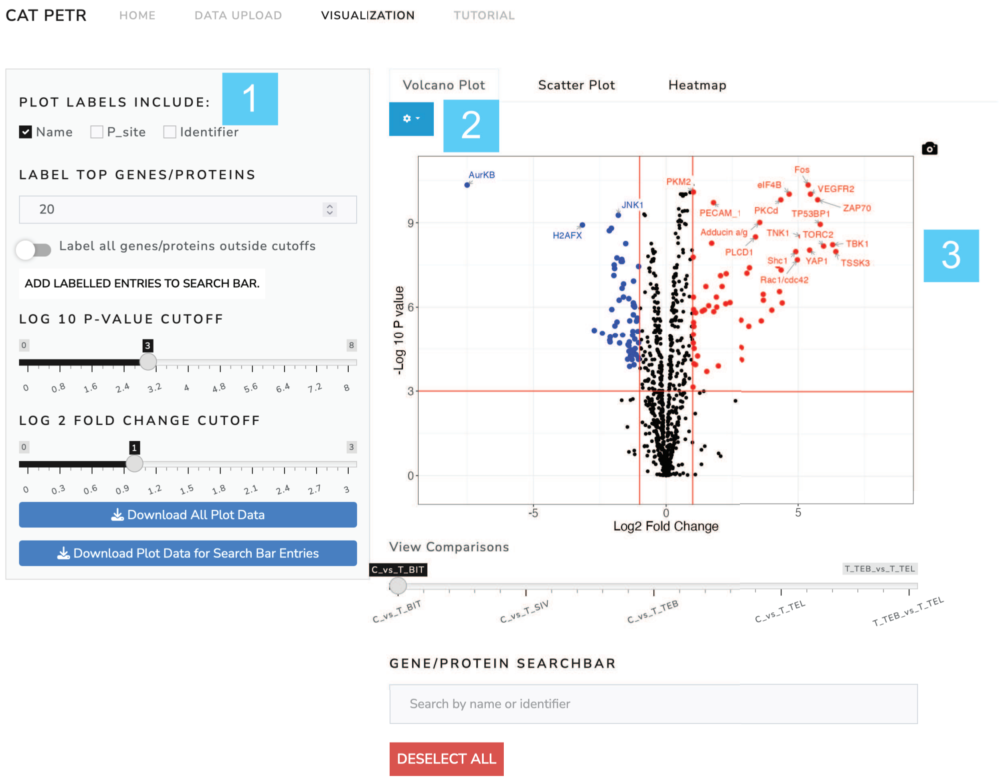

Figure 4: All three of the visualization sub tabs can be broken into three sections: (1) A sidepanel with various inputs that alter the plot parameters and buttons to download the plot data. (2) an appearance drop down menu which allows for superficial alterations to the appearance of the plots. (3) A main panel which contains the plot itself and a gene/protein search bar that allows for specific genes to be labelled on the plot.

8. Use the sidebar to alter the parameters of the volcano plot in the following ways:
   (a) Add the P_sites to the volcano plot labels using the checkboxes.
   (b) Reduce the number of top labelled proteins to 10.
   (c) Adjust the log10 P-value cutoff to 2 and the log2 fold change cutoff to 1.5.
9. Use the appearance option menu to alter the appearance of the volcano plot in the following ways:
   (a) Increase the size of the plot to 760 width by 590 height.
   (b) Increase the label text size to 5.
   (c) Change the color of the searchbar points.
10. Highlight specific genes/proteins using the protein search bar in the following ways:

(a) Begin typing the protein name 'PKCg T655 PK083' into the search bar and then select it from the resulting drop down menu. Note that the protein becomes labelled in green on the plot. Select 'PKCg T655 PK083' on the search bar by clicking on it and then remove it using the delete key.

(b) Click on any random point on the graph to add it to the search bar and generate a label. Then click on the point again to remove it.

(c) Click the 'add labelled entries to search bar' button. Note that the color of the labels have all switched to the color we specified in step 9.

11. Switch to the 'C_vs_T_SIV' plot using the slider directly underneath the plot.

12. Download a .svg image of the plot by clicking on the small camera icon displayed on the upper right hand corner of the plot.

13. The buttons at the bottom of the sidebar allow the user to either download all of the data for the currently viewed volcano plot, or just the data from genes/proteins that are currently in the sidebar. For now, hit the button that downloads just the search bar data and save the resulting tsv file on your computer.

##### 3.2.2 Scatter Plot

The scatter plot tab works very similarly to the volcano plot tab in many ways. As such, the details of downloading the plot and its data, the contents of the sidebar and appearance menu, and how to add/remove proteins from the search bar will not be covered in this tutorial.

The major difference of the Scatter plot section is that in order to view the scatter plot it is necessary for the user to define which axes will show the fold changes from which comparison.

14. Switch to the scatter plot sub-tab. There should be no visible plot and the sub-tab will have the appearance shown below.

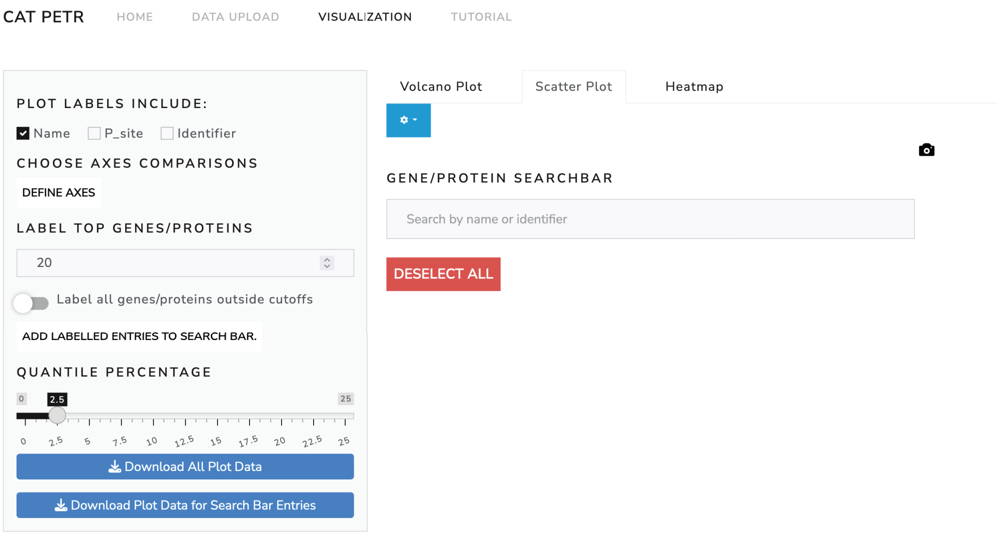

Figure 5: Blank scatter plot page.

15. Hit the 'define axes' button to bring up a modal which, similarly to step 5, brings up 3 bucket lists (see image below). Just like in step 5, we can drag our comparisons over to the X and Y-axis boxes to determine what data will be shown on the graphs. Drag and drop the comparisons to match the image below and then hit the generate plots button.

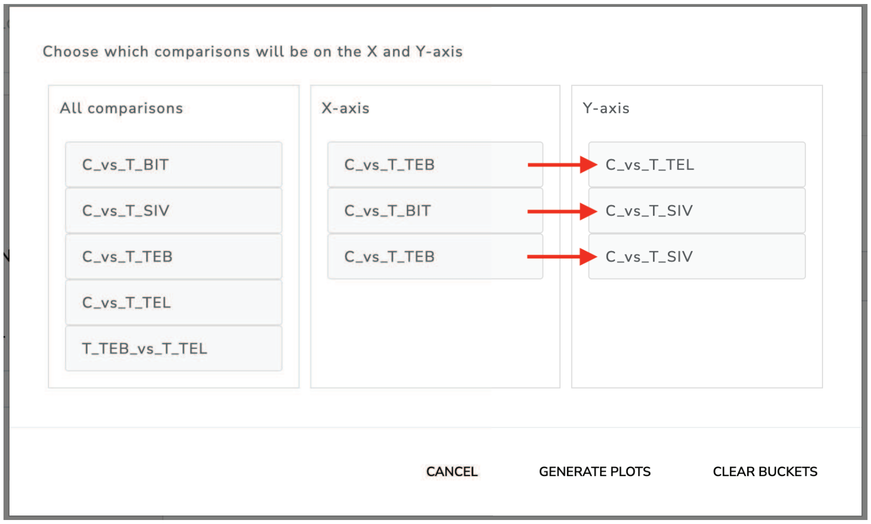

Figure 6: Example X and Y-axis selection bucket list

16. Hit the deselect all button below the gene/protein searchbar in order to clear it.

17. Click the 'add labelled entries to search bar' button.

##### 3.2.3 Heat map

The Heatmap sub tab is unique in that the contents of the plot are entirely dependent on the contents of the gene/protein searchbar. If the gene/protein searchbar is empty, then no plot will appear. The contents of the searchbar should be full from step 17.

18. Switch to the heatmap sub-tab. The heatmap should load automatically and should have the appearance shown below.

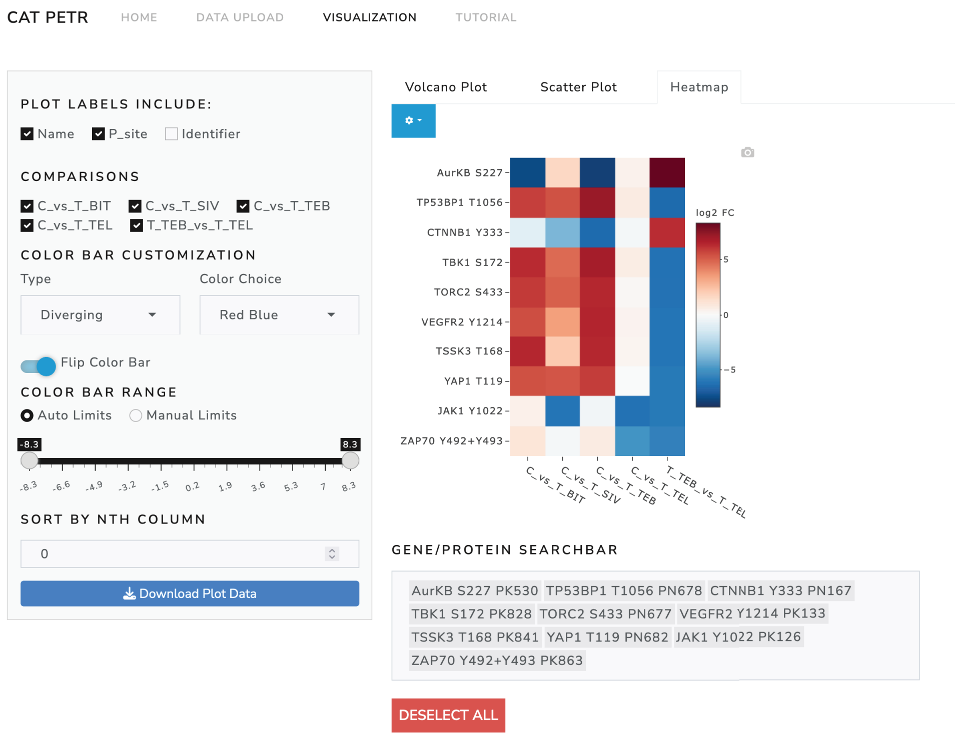

Figure 7: Example heatmap page.

19. Mouse over the heatmap to observe the information popups that appear for each tile.

20. Use the sidebar to alter the parameters of the heatmap in the following ways:

    (a) Remove the 'T_TEB_vs_T_TEL' comparison from the heatmap by un-checking it.

    (b) Change the color bar color choice to "Red Yellow Blue".

    (c) Adjust the colorbar range to -7 and 7.

    (d) Enter 1 into the 'sort by nth column' input to sort the heatmap by the first column.

21. Use the appearance option menu to alter the appearance of the heatmap in the following ways:

(a) Decrease the label size to 11.
(b) Decrease the tile width to 35.
(c) Decrease the tile height to 20.

22. Save the plot as a .svg using the small camera icon on the upper right corner of the heatmap.

23. Download the plot data using the button at the bottom of the sidebar.

### 4 Appendix

#### 4.1 Normalization and VSN

In essentially all forms of high-throughput expression/phosphorylation data there is a relationship where the variance of the intensities is proportional to the mean of the intensities. This fundamentally breaks the assumption of equal variance that is required for many statistical tests, including t-tests. Therefore, normalization is often used to stabilize the variance across different intensities. ( [3]). CAT PETR offers two different methods for normalization. The first is a log transformation. Log transformation is a simple and widely used method for controlling variance, but it comes with certain drawbacks. For example, log transformation is unable to handle negative values, which often occur in microarray data due to background correction. A more advanced method that addresses the issues associated with log transformation is variance stabilization normalization (VSN) ( [2]). VSN uses an $arsinh$ based transformation with parameters optimized via a robust variant of maximum-likelihood estimation. This optimization works on the assumption that a majority of the genes/proteins tested have no change in expression/phosphorylation; an assumption that is valid for the vast majority of expression/phosphorylation studies. Unlike log transformation, VSN is able to handle intensities at or below zero. In addition, VSN generally gives more accurate variance stabilization for near zero intensities, which tend to have their variances inflated by log transformation.

#### 4.2 Cyber-T

The collection of high-throughput expression and phosphorylation data is a difficult and costly process. Therefore, many studies have very limited sample sizes which greatly limits the statistical power of any differential analysis. This problem is compounded by the fact that extensive multiple testing correction must be completed due to the high number of genes/proteins being tested. As such, methods that can adjust for limited sample sizes and allow for more statistical power have become a fundamental part of differential expression/phosphorylation analysis. CAT PETR uses one such method called the CyberT method. CyberT uses the weighted average of the empirical variance and a prior variance estimate to obtain a more accurate estimation of variance within each treatment group ( [4]). The empirical variance is measured using the difference in intensity between biological replicates. The prior estimate is derived by pooling the variances of genes/proteins in the same neighborhood (e.g. genes/proteins with similar intensities). The size of this neighborhood is controlled by a sliding window. The size of the sliding window and the amount of weight (confidence) placed on the pooled variance estimate should be adjusted based on the number of genes/proteins tested and the number of biological replicates respectively. CAT PETR automatically chooses the optimal values for the sliding window and confidence based on the recommendations from [1] This optimization means that the effect of this pooled variance will decrease with an increasing number of biological replicates. For more information on Cyber-T, please see [1] and [4].

#### 4.3 Multiple Testing Correction

Regardless of whether a traditional student's t-test or a Cyber-T t-test is used for the analysis, the P-values are corrected via the Benjamini-Hochberg method. ( [5]).
